# Excitation and inhibition delays within a feedforward inhibitory pathway modulate cerebellar Purkinje cell output in mice

**DOI:** 10.1101/2023.01.18.524515

**Authors:** Francesca Binda, Ludovic Spaeth, Arvind Kumar, Philippe Isope

**Author notes:** These authors contributed equally (Francesca Binda, Ludovic Spaeth). Francesca Binda: Dept. of Fundamental Neurosciences, University of Lausanne, CH-1005 Lausanne, Switzerland. Ludovic Spaeth: Dominick P Purpura Department of Neuroscience, Albert Einstein College of Medicine, Bronx, New York, NY 10461, United States of America.

## Abstract

The cerebellar cortex computes sensorimotor information from many brain areas through a feedforward inhibitory (FFI) microcircuit between the input stage, the granule cell layer, and the output stage, the Purkinje cells. While in other brain areas FFI underlies a precise excitation vs inhibition temporal correlation, recent findings in the cerebellum highlighted more complex behaviors at the granule cell (GC) – molecular layer interneuron (MLI) – Purkinje cell (PC) FFI pathway. To dissect the temporal organization of the cerebellar FFI pathway, we combined ex *viv*o patch clamp recordings of PCs with a viral-based strategy to express Channelrhodopsin2 in a subset of mossy fibers (MFs), a major excitatory input to GCs. We show that light-mediated MF activation elicits excitatory and inhibitory currents in PCs with a wide range of temporal delays. Furthermore, in many recordings, excitation and inhibition were initiated by different groups of GCs, expanding PCs synaptic temporal integration. Using a computational model of the FFI pathway we demonstrated that this temporal expansion could strongly influence how PCs integrate MF inputs. Our findings suggest that MF inputs are also encoded by specific delays between excitation and inhibition in PCs.

## Introduction

In the cerebellar cortex, GCs and MLIs define a feedforward inhibitory (FFI) pathway triggering in PCs a reliable excitation/inhibition sequence (E/I) with a very low temporal dispersion (Eccles et al., 1967; Brunel et al., 2004; Mittmann et al., 2005; Grangeray-Vilmint et al., 2018, **Figure 1A, first panel**). This fast sequence is favored by the beam-like arrangement of parallel fibers, the axon of the GCs, which run perpendicularly to MLIs and PCs dendrites in the mediolateral axis. Moreover, MLI axons target essentially neighboring PCs in the sagittal plane (Palay and Chan-Palay, 1974; Jörntell et al., 2010; Kim and Augustine, 2021). Parasagittal arrays of neighboring PCs constitute discrete computational units called microzones, the cortical part of cerebellar modules (Apps and Hawkes, 2009; Apps et al., 2018). Therefore, an appealing hypothesis is that each GC axon contacts both MLI and PC dendrites within a given cerebellar microzone leading to the modulation of PC discharge, a feature that we will define as the classical FFI. FFI sharpens the temporal window for spike discharge, prevents PC discharge saturation and promotes long-term plasticity at GC-PC synapses, thus extending the dynamic range for GC inputs encoding (Eccles et al., 1967; Barbour, 1993; Brunel et al., 2004; Mittmann et al., 2005; Isaacson and Scanziani, 2011; Binda et al., 2016; Rowan et al., 2018). However, most of these studies were performed by stimulating either bulks of parallel fibers or GC somas, while in natural conditions FFI is rather triggered by mossy fibers (MF) exciting more discrete, spatially organized, clusters of GCs.

**Figure 1.**
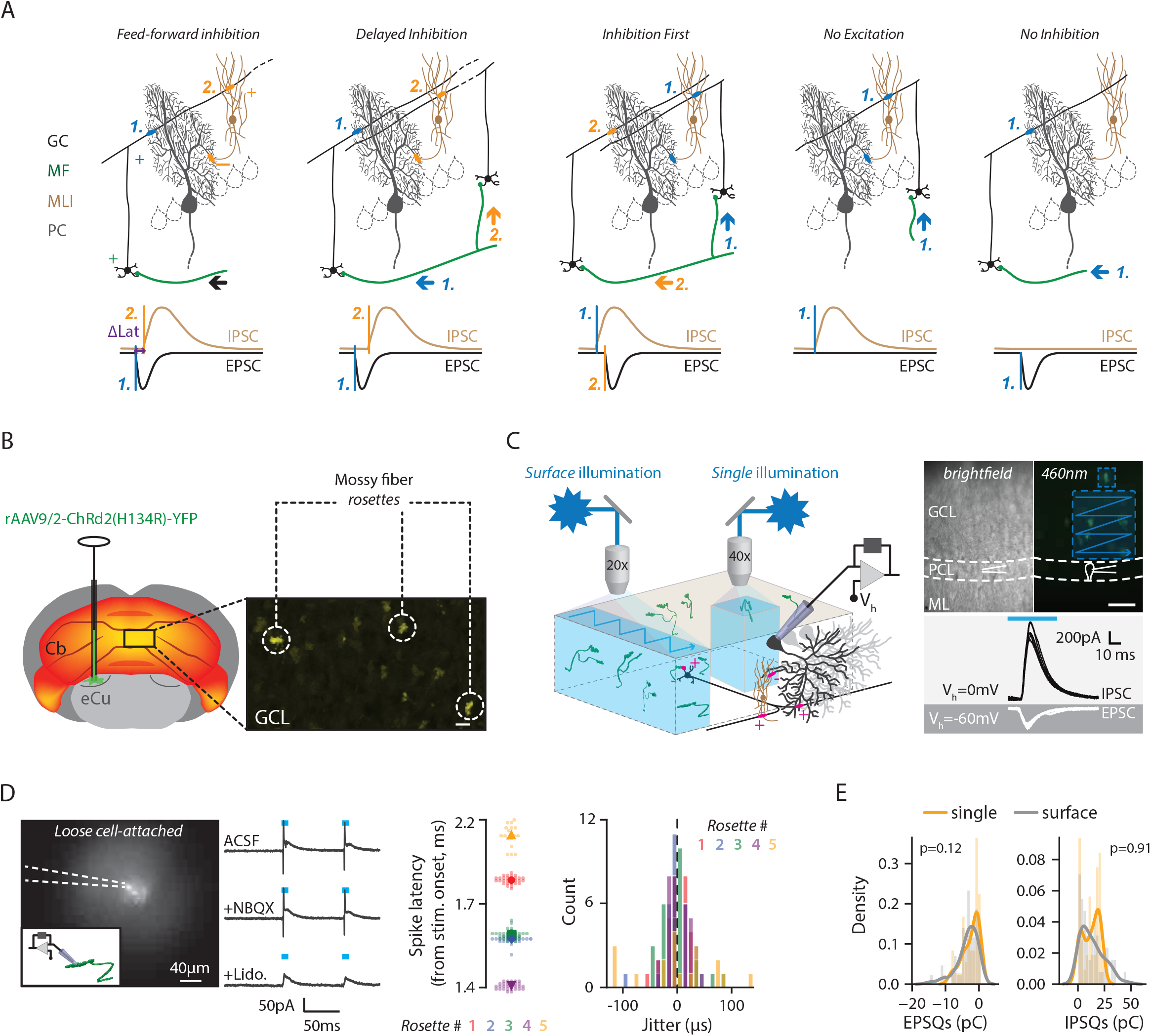
MF optogenetic stimulation elicits EPSC and IPSCs in PCs. **A**, diagram of the cerebellar cortex illustrating how MF inputs affect the timing of GC-PC and GC-MLI-PC inputs. GC clusters targeted by a given MF (in green) may lead to independent excitatory (EPSC) and inhibitory (IPSC) synaptic inputs, therefore yielding different types of E/I sequences in PCs: (1) a classical FFI with a short E/I delay (ΔLat = latency of inhibition – latency of excitation, Feed-forward inhibition panel); (2) A long or a reverse ΔLat when several GC clusters lead to independent E/I sequence (Inhibition delayed or inhibition first panels); (3) no E/I sequence (no excitation and no inhibition panels). **B**, *Left*, rAAVs 9/2 injection in the external cuneate nucleus allowing further expression of ChannelRhodopsin-2-YFP in cerebellar mossy fibers. *Right*, confocal image of YFP fluorescence in MF rosettes from a transverse cerebellar slice. **C**, *Left*, protocols of photostimulation. In surface illumination, several MF terminals are activated sequentially in an 83×83 μm area. In single illumination, a patterned light beam is selectively applied to a unitary rosette. *Top right*, example of acute slices in situ, under IR or 460 nm light illumination. The large and small dashed-blue squares illustrate surface and single illumination, respectively. Scale bar 40 microns. *Bottom right*, example of inhibitory (IPSC, Vh = 0mV) and excitatory (EPSC, Vh = −60mV) post synaptic currents elicited in PCs and evoked by MF photostimulation. GCL: granule cell layer, PCL: Purkinje cell layer, ML: Molecular layer. **D**, *Left*, example of loose cell-attached recording of a rosette and corresponding current evoked by photostimulation in normal ACSF (see methods), in presence of NBQX and NBQX + Lidocaine (10 μM each). *Middle*, latency of the first spike from stimulation onset elicited in 5 different rosettes. Large and small markers show average and individual trials respectively. *Right*, variability in spike offset (jitter) measured in rosettes. The latency of each spike in every trial was subtracted to the average latency of all the trials (each color represents a different rosette). **E**. Distribution of synaptic charges measured in EPSCs and IPSCs evoked by photosimulation, in both surface (light grey) and single (orange) illumination protocols. EPSQ, U=1432.0; IPSQ, U=1199.0; Mann Whitney U test, p-values in plot.

MFs, which originate in pre-cerebellar nuclei in the brainstem and spinal cord, make excitatory synapses onto GCs and convey sensorimotor, proprioceptive and vestibular information to the cerebellar input stage. A single MF gives rise to numerous collaterals and contact many GC clusters in a given lobule (Shinoda et al., 2000; Quy et al., 2011). This arrangement leads to temporally dispersed GC activation as observed in lobule IX and X of the cerebellar cortex (Chabrol et al., 2015). Furthermore, clusters of GCs make active synaptic connections with specific groups of PC and MLIs independently as a result of activity-dependent processes (Barbour, 1993; Isope and Barbour, 2002; Valera et al., 2016; Spaeth et al., 2022). Indeed, PCs and MLIs located in the same cerebellar module have specific GC input connectivity maps, suggesting that GCs may contact either MLIs or PCs at a given location. *In vivo* recordings confirmed that sensory inputs elicit multiple temporal combinations of excitation and inhibition in PCs (Santamaria et al., 2007; Jelitai et al., 2016; Brown and Raman, 2018). These combinations include classical FFI sequences, but also inhibition preceding excitation or inhibition without excitation, supporting the hypothesis that PCs may combine separate excitatory and inhibitory synaptic inputs originating from spatially distinct GCs clusters. Therefore, we postulate that sequences of excitation (E) and inhibition (I) in the FFI pathway may have a wide range of delays when reaching PCs (**Figure 1A**). We therefore explored this hypothesis by combining optogenetic MF stimulation and *ex-vivo* cerebellar recordings. Channelrhodopsin2 was expressed in a subset of MFs terminals allowing selective MF activation and subsequent measure of E/I delays in PCs with whole-cell voltage-clamp. We provide a direct demonstration that the GC-(MLI)-PC FFI undergoes temporal expansion, beyond canonic FFI. Finally, using a computational approach, we suggest that this arrangement expands PC coding capacity, which would further increase their ability to process and integrate multiple sensorimotor inputs, strengthening the ability of the cerebellum to contribute to the performance of various and diverse tasks and behaviors in many different contexts.

## Results

### Stimulating specific MFs and individual rosettes

In order to physiologically recruit the GC-(MLI)-PC FFI pathway in the cerebellar cortex, we selectively expressed Channelrhodopsin2 in individual MFs by injecting a rAAV 9/2-hSynChRd2(H134R)-YFP in the external cuneate nucleus (eCu), a pre-cerebellar nucleus (**Figure 1B**; number or recordings n= 113, number of cells n = 24 and number of mice N = 8), which convey proprioceptive and exteroceptive information from the upper limbs. MFs originating in the eCu send collaterals to different location in a given lobule (Quy et al., 2011; Valera et al., 2016). We then photostimulated MFs in acute cerebellar slices while recording vermal PCs (lobule III to VIII) using whole-cell patch-clamp and monitored synaptic input current from the FFI pathway (**Figure 1A, 1C**, see methods). Two strategies of photo stimulation were developed: (1) a laser scanning method in order to illuminate small surfaces of the GC layer (83 × 83 μm, hereinafter referred to as *surface illumination*) leading to the activation of many MF rosettes (**Figure 1C**). This method mimics the desynchronized activation of several MFs. (2) alternatively, we used patterned illumination (see Methods) to specifically illuminate a single rosette and activate one MF (hereinafter referred to as *single illumination*, **Figure 1C**). We first addressed whether blue light enables individual MF rosette excitation. YFP fluorescence allowed us to visualize and record MF rosettes in loose cell-attached mode (Barbour and Isope, 2000; **Figure 1D**). In all the rosettes, illumination triggered a direct depolarization eliciting a unique and reliable action potential (mean delay± SD: 1.63 ± 0.31 ms; mean jitter ± SD: 0.023 ± 0.03 ms; n=5; **Figure 1D**). Application of NBQX and Lidocaine isolated the direct depolarizing current elicited during the length of the light pulse (mean amplitude ± SD, 15.6 +/-9.8 pA, n=5, **Figure 1D**). Therefore, illumination likely releases a major part of the full ready releasable pool of vesicles at the MF-GC synapses ensuring reliable activation of connected GCs.

### MF illumination elicited temporally dispersed E and I with correlated synaptic weights

We recorded excitatory and inhibitory post synaptic currents (EPSCs and IPSCs) in PCs upon MF activation via single or surface illumination. EPSCs and IPSCs were recorded at V_m_ = −60 mV and V_m_ = 0 mV respectively (**Figure 1C**). To estimate the total synaptic weight transfer, we calculated the charge (excitatory - EPSQ and inhibitory – IPSQ) carried by these currents (mean ± SD, surface illumination, EPSQ: - 3.4 ± 3.1 pC; IPSQ, 13.4 ± 10.8 pC, n=54 recordings, n = 17 cells; single illumination: EPSQ: −2.5 ± 2.5 pC; IPSQ: 12.3 ± 7.6 pC, n= 45 recordings, n= 7 cells; **Figure 1E**). MF photostimulation elicited both excitation and inhibition in PCs at varying delays (surface illumination: mean delay excitation ±SD: 14.1 ± 23.6 ms; mean delay inhibition ±SD: 18.1 ± 4.3ms; single rosette: mean delay excitation ± SD: 5.6 ± 1.5 ms; mean delay inhibition ± SD: 7.4 ± 2.0 ms). The delay observed in excitatory currents recorded in PCs includes the sum of (1) the time for MF rosette depolarization and synaptic release, (2) the integration time in GCs, (3) the conduction time to GC synaptic boutons and finally (4) the synaptic release time to PC dendrites. The delay observed in inhibitory currents includes one additional synapse and integration time in MLIs. Since MLIs contact only neighboring PCs (Palay and Chan-Palay, 1974; Kim et al., 2014), if a given group of GC axons contacts both MLIs and PCs, the delay between excitation and inhibition on PC dendrites should be short and invariant (first scenario in **Figure 1A**). However, using blue light illumination of MFs or individual rosettes, we observed a large temporal spread of the delay between excitatory and inhibitory synaptic inputs recorded in PCs named ΔLat (i.e. IPSC latency – EPSC latency). In both types of illumination, ΔLat ranged from negative to positive values that can exceed ± 5ms (*surface illumination*: from −3.14 to 21 ms, mean ± SD = 3.9 ± 4.0 ms, n=54; *single illumination*: from −5.9 ms to 4.2 ms, mean ± SD = 1.75 ± 1.78 ms, n=44 **Figure 2A, 2B**). These results indicate that even when stimulating a single rosette yielding inputs from a single MF, several independent GC-PC and GC-MLI-PC pathways are activated, as observed by the presence of specific ΔLat.

**Figure 2.**
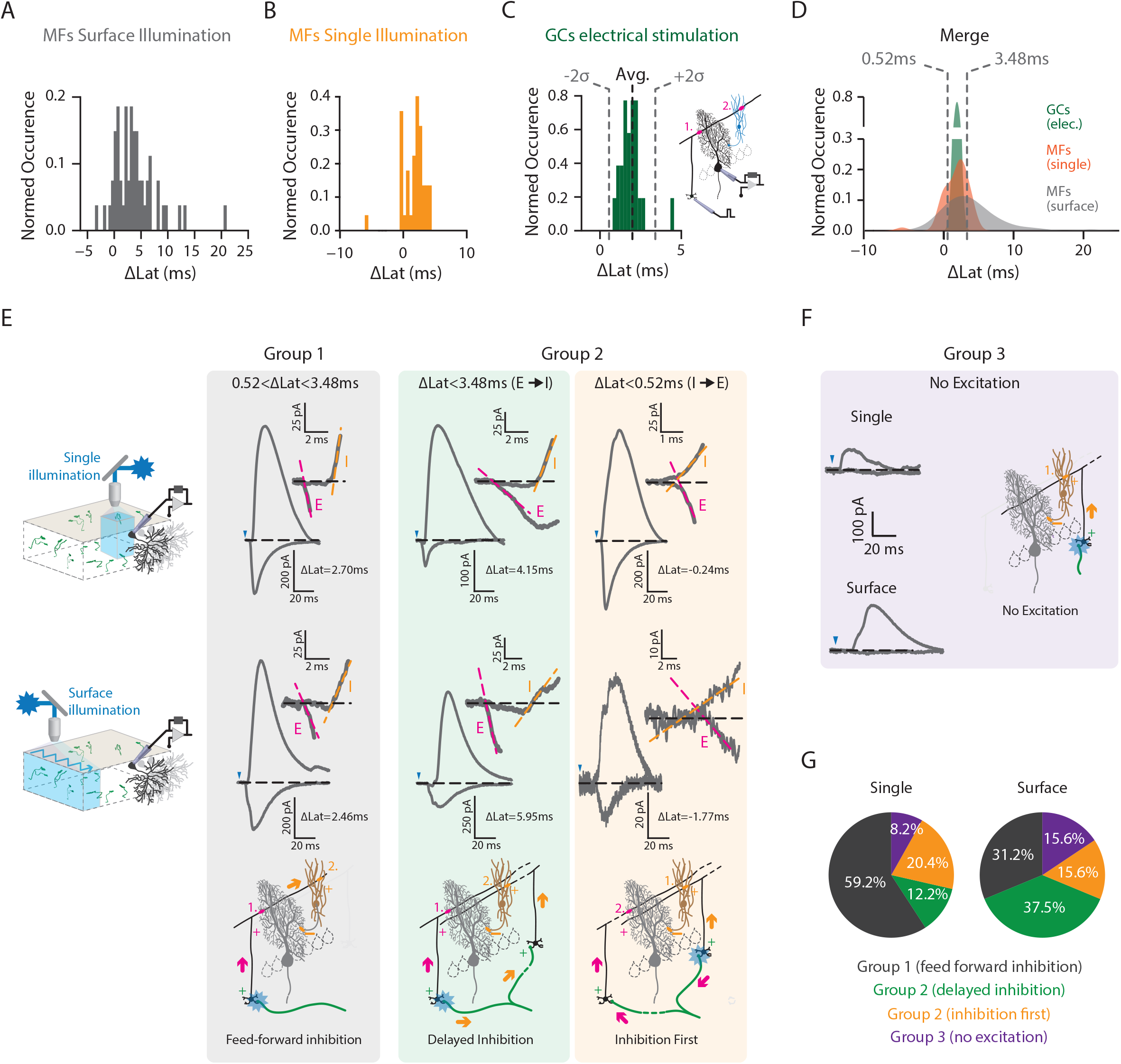
MF stimulation elicits a wide range of ΔLat profiles in PCs. **A**, histogram of the ΔLat following MF surface illumination. **B**, histogram of the ΔLat following single rosette illumination. **C**, histogram of the ΔLat following GC electrical stimulation (direct FFI, GC-ΔLat). The dotted black line represents the average and gray lines illustrate the interval containing ΔLat considered as direct FFI (i.e., mean±2 SD: 0.52<Δ Lat<3.48ms). Inset, diagram illustrating FFI obtained with direct excitation of GCs. **D**, Merged ΔLat distributions from MF surface illumination, single rosette illumination and GCs electrical stimulation. Data between dotted lines illustrate ΔLat considered as direct FFI. **E**, representative examples of E/I temporal profiles observed during single rosette and surface illumination. Group 1 corresponds to 0.52 < ΔLat < 3.48ms, illustrating direct FFI. Group 2 correspond to larger (Inhibition delayed) or negative (inhibition first) ΔLat. Blue triangles represent stimulation onset. Inset, onset of EPSC and IPSC superimposed. Magenta and orange dashed lines show linear fit of current onsets. Latency was measured as the intersection with baseline (black dashed line). **F**, in a fraction of experiments, no excitation could be recorded following photostimulation (Group 3). This illustrates GC clusters eliciting only inhibition. Excitation only was never observed in our conditions. **G**, proportion of the different scenarios observed in all recordings from single and surface illumination protocols.

As control, we stimulated individual groups of GCs using electrical stimulation (**Figure 2C**, see methods). We then observed short and almost invariant E/I ΔLat (mean GC_ΔLat = 2 ± 0.74 ms, mean ± SD, n = 23; **Figure 2C**) as previously shown (Vincent and Marty, 1996; Brunel et al., 2004; Mittmann et al., 2005; Grangeray-Vilmint et al., 2018). These reliable ΔLat accounted for classical FFI (**Figure 1A**). We then used these GC_ΔLat as a proxy for the classical FFI in the GC-MLI-PC pathway. Following MF stimulation, we classified an E/I sequence as a classical FFI when ΔLat exceeded GC_ΔLat ± 2 SDs (i.e. 0.52 ms < ΔLat < 3.48 ms, referred as group 1; group 2 correspond to indirect FFI; in group 3, only inhibition was recorded, **Figure 2D-F**). Therefore, we identified that E/I balance in PC somas is frequently determined by independent clusters of GCs (surface illumination: group 2/3 = 68.8 %; single illumination: group 2/3 = 40.8 %; **Figure 2G**).

In the forebrain, excitation-inhibition (E/I) ratios are equalized (i.e., inhibitory inputs are scaled according to their excitatory inputs, Xue et al., 2014). We then evaluated whether excitatory and inhibitory synaptic charges elicited by MF stimulation are correlated. In both classical FFI (group 1) and other sequences (group 2), the excitatory synaptic charges (EPSQs) were linearly correlated with the inhibitory synaptic charges (IPSQs; **Figure 3A-C**). Thus, in GCs targeted by the same MF, the excitatory and inhibitory synaptic weights evoked in PCs are paired together and lead to homogeneous and scaled E/I balance as described in the cerebral cortex (E/I in single protocol, 0.2 ± 0.12, n = 45; in surface protocol, 0.46 ± 0.79, n = 54; mean ± SD, **Figure 3D**, Xue et al., 2014). In agreement with these results, we found that inhibition was significantly stronger when MF stimulation led to both excitatory and inhibitory inputs rather than when only inhibition was measured (**Figure 3E**).

**Figure 3.**
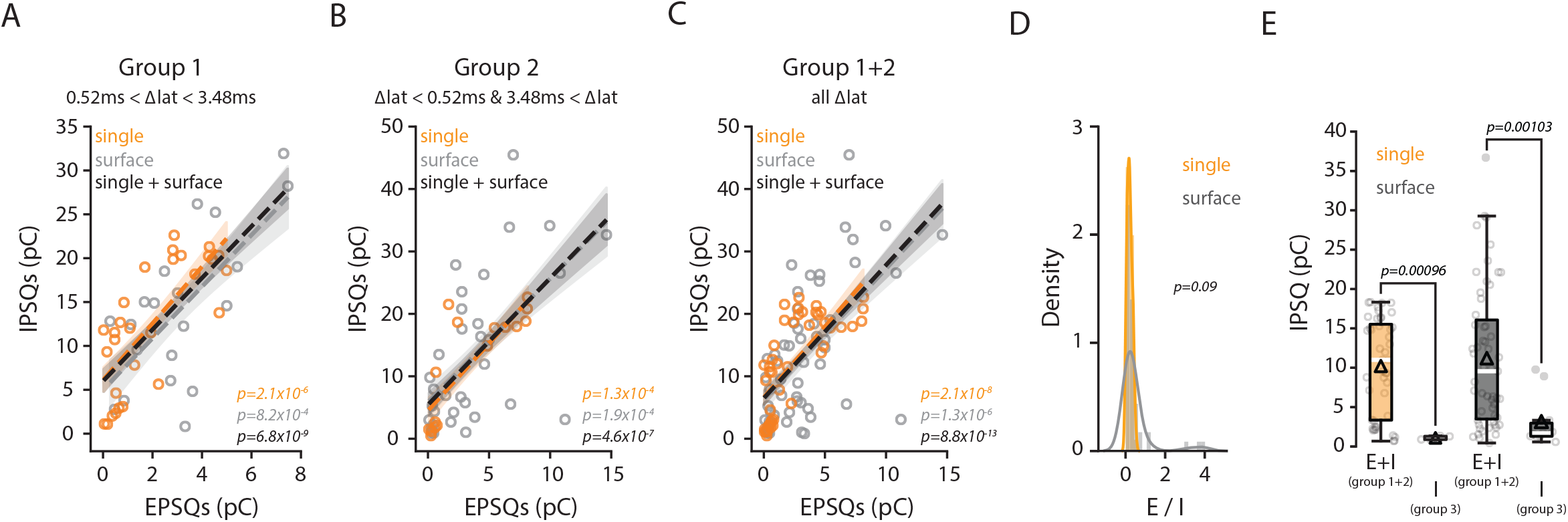
E/I synaptic weights are correlated. **A**, scatter plot of inhibitory synaptic charges (IPSQ) against excitatory synaptic charges (EPSQ) from E/I sequences considered as FFI (Group 1). Orange, single illumination (r=0.76, n=29, p-value in plot); light gray, surface illumination (r=0.68, n=20, p-value in plot); black, linear fit of merged datasets (r=0.72, n=49, p-value in plot). Colored interval represents 95%CI in all cases. **B**, scatter plot of inhibitory synaptic charges (IPSQ) against excitatory synaptic charges (EPSQ) from group 2 E/I sequences. Orange, single illumination (r=0.81, n=16, p-value in plot); light gray, surface illumination (r=0.59, n=34, p-value in plot); black, linear fit of merged datasets (r=0.64, n=50, p-value in plot). Colored interval represents 95%CI in all cases. **C**, scatter plot of inhibitory synaptic charges (IPSQ) against excitatory synaptic charges (EPSQ) from all E/I sequence (Group 1 + 2). Orange, single illumination (r=0.72, n=45, p-value in plot); light gray, surface illumination (r=0.6, n=54, p-value in plot); black, linear fit of merged datasets (r=0.64, n=99, p-value in plot). Colored interval represents 95%CI in all cases. **D**, Histogram of the synaptic charge E/I ratio (orange, single; gray, surface illumination). Curves show kernel density estimation of distributions. Mann-Whitney test, U=970.0, p-value in plot. **E**, Comparison between inhibitory synaptic charges from group 3 vs group 1+2. Black triangle, average; black line, median; box edges, interquartile range; whiskers, min & max; filled circles, outliers. Single, E+I (n=44) vs I Only (n=4), U=166.0. Surface, E+I (n=55) vs I Only (n=10), U=456.0; Mann-Whitney test in both, p-values in plot.

### Variable delays expand PC dynamics

We anticipated that this broad distribution of ΔLat in PC soma might influence PC discharge. Since optogenetic MF illumination at high frequencies was not possible, we used a computational model of the GC-MLI-PC FFI pathway developed in a previous study (Grangeray-Vilmint et al., 2018) (see Methods) and assessed how ΔLat affects PC discharge for various synaptic weights including presynaptic short term dynamics parameters (i.e. at GC-MLI and MLI-PC connections, STD) and stimulation frequencies. In this model, excitatory post synaptic potentials (EPSPs) and inhibitory post synaptic potentials (IPSPs) amplitudes as well as STD at excitatory and inhibitory synapses were simulated using the deterministic model proposed by Tsodyks et al. (Tsodyks and Markram, 1997; see **Table 1**) in which a single parameter (*U*) can be used to systematically change the nature of a synapse from a facilitating to a depressing mode. Four parameters described the E/I balance (EPSP and IPSP amplitudes) and the STD of excitation (U_E_) and inhibition (U_I_). In our previous study (Grangeray-Vilmint et al., 2018) we demonstrated that these four parameters in association with the number of GC burst stimulations strongly influence how GCs control PC discharge. Since, on average, GC-PC synapses facilitate while GC-MLI-PC synapses depress (Grangeray-Vilmint et al., 2018), GC bursts (3 stimulations) leading to a net inhibition can switch to a net excitation when burst duration increases (e.g. 7 stimulations). The GC-MLI-PC FFI pathway model with short-term dynamics can reproduce PC discharge following burst of GC inputs (3, 5 and 7 stimulations) in various combination of parameters. We therefore quantified changes in PC firing rate spike gain (see Methods). The spike gain is negative when GC stimulation led to omitted spikes (i.e net inhibition) during the time window of the stimulation and positive when an excess of spike (i.e. net excitation) is simulated.

**TABLE 1.** Model parameters.

| **Neuron Parameters** |  |
| --- | --- |
| Membrane Capacitance ($C_m$) | 250 pF |
| Membrane time constant ($\tau_m$) | 20 ms |
| Resting membrane potential ($V_{rest}$) | -70 mV |
| Refractory period ($\tau_{ref}$) | 2 ms |
| Spike threshold ($V_{th}$) | -55 mV |
| **Excitatory Synapse Properties** |  |
| Excitatory reversal potential | 0 mV |
| Excitatory time constant | 1.0 ms |
| Static unitary EPSP | 1.22, 1.52, 1.82 mV @ -70 mV |
| **Short-term dynamics of excitatory synapses** |  |
| Synaptic decay time constant ($\tau_{psc}^{exc}$) | 1.5 ms |
| Recovery time constant ($\tau_{rec}^{exc}$) | 30 ms |
| Facilitation time constant ($\tau_{fac}^{exc}$) | 500 ms |
| Fraction of opened Calcium channels ($U^{exc}$) | 0.03 or 0.05 |
| Synaptic amplitude scaling ($A^{exc}$) | $A^{exc} = A_e / U^{exc}$ where $A_e = \{2.0, 2.5, 3.0\}$. This resulted in unitary EPSP amplitude of 1.22, 1.52, 1.82mV at resting membrane potential |
| **Inhibitory Synapse Properties** |  |
| Inhibitory reversal potential | -80 mV |
| Inhibitory time constant | 5 ms |
| Static unitary IPSP | {-0.63, -1.09, -1.39 mV} @ -70 mV |
| **Short-term dynamics of inhibitory synapses** |  |
| Synaptic decay time constant ($\tau_{psc}^{inh}$) | 1.5 ms |
| Recovery time constant ($\tau_{psc}^{inh}$) | 100 ms |
| Facilitation time constant ($\tau_{psc}^{inh}$) | 800 ms |
| Fraction of opened Calcium channels ($U^{inh}$) | 0.3 |
| Synaptic amplitude scaling ($A^{inh}$) | $A^{inh} = A_i / U^{inh}$, where $A_i = \{1.5, 2.5, 3.5\}$. This resulted in unitary IPSP amplitude of -0.63, -1.09, -1.39mV) |
| **Excitation inhibition (E/I) delay** |  |
| EI delay | -5ms (I before E) to +6ms (E before I) |

Using biologically plausible range of EPSP (1.22 mV to 1.82 mV at resting potential), IPSP (−0.63 mV to −1.39 mV at resting potential), STD parameters (facilitation at GC-PC synapses, depression at MLI-PC synapses) and stimulation frequencies (3-7 stimulations at 10-200 Hz) (Chadderton et al., 2004; Jörntell and Ekerot, 2006; Arenz et al., 2008; Grangeray-Vilmint et al., 2018), we could reproduce sample raster plot observed in biological data (**Figure 4A** raster plots; see Grangeray-Vilmint et al., 2018;). Next, we estimated spike gain by systematically varying ΔLat from −5 ms (I before E) to +6 ms (E before I), input frequency (10-200 Hz), and stimulus burst size (3,5,7) for different sets of E, I, U_E_ and U_I_ (**Figure 4A,B**, Table 1; see also Materials and Methods). **Figure 4C and D** show spike gains for the same set of E, I, U_E_ and U_I_ parameters as a function of ΔLat and stimulation frequency for different burst durations.

**Figure 4:**
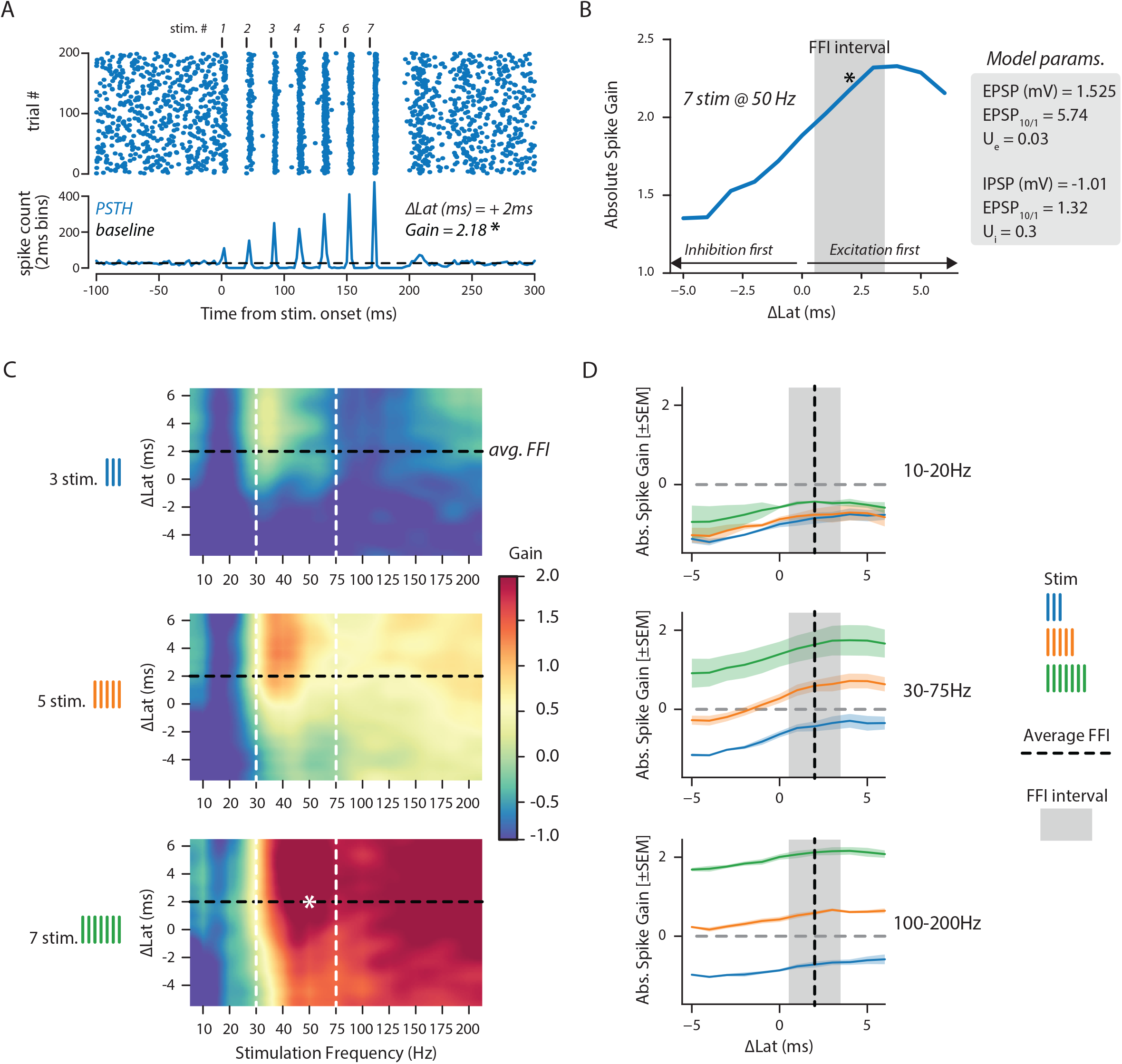
Spike gain in PCs relies on ΔLat and stimulation frequency. **A**, Illustrative raster plot (top) and peristimulus time histogram (bottom) simulating PC discharge and spike gain (i.e., mean normalized PSTH) in response to 7 stimulations delivered at 50Hz with a fixed ΔLat of +2ms. **B**, absolute spike gain as a function of various ΔLat (blue curve) with input parameters descried in the gray box. The gray box illustrates the range of classical FFI. The black asterisk shows the gain measured in the example data shown in A. **C**, spike gain as a function of variable ΔLat (−5ms to +6ms) and stimulation frequencies (10 Hz to 200 Hz). **D**, average spike gain measured across 3 (blue curves), 5 (orange curves) or 7 (green curves) stimulations delivered at frequencies ranging from 10 to 20Hz (top), 30 to 75Hz (middle) and 100 to 200Hz (bottom). The black vertical dashed line represents the average ΔLat found in FFI. The gray box shows the FFI interval (+/-2SD from average).

We observed that while spike gain is negative for short burst of GC stimulation (top left of Figure 4C), it becomes positive for longer burst (left bottom of Figure 4C) at all frequencies above 30 Hz and for most ΔLat. Indeed, for short bursts, when inhibition comes a few milliseconds before the excitation the impact of EPSP on the spike probability of the neuron is strongly reduced (blue part in the upper left panel, **Figure 4C**). By contrast, when inhibition comes a few milliseconds after the excitation, the spike probability may not be much affected as the neuron may have already spiked when the EPSP peaked, i.e., before peak of the incoming IPSP. Consistent with this reasoning, as we increase the ΔLat from negative to positive, the spike gain increased monotonically with a steady state above ΔLat = 2.5 ms (vertical axis of left panels in **Figure 4B, C**). However, this absolute spike gain increase was strongly affected by the GC stimulation frequency. This is because excitatory and inhibitory synapses undergo STD (Grangeray-Vilmint et al., 2018) and the amount of facilitation and depression per spike differ for excitatory and inhibitory synapses. These simulations also revealed that ΔLat influence is maximal at stimulation frequency of 30-75Hz as also observed when spike gains were pooled by stimulus frequency bands (**Figure 4C-D**,). Overall, similar patterns (reduced spike probability when inhibition comes first and monotonic increase in absolute spike gain) were observed when the stimulus consisted of 5 or 7 spikes. (**Fig 4C**, Left middle and bottom). For longer stimuli and higher frequencies, the definition of the ΔLat becomes blurred from second to the last spike and the influence of ΔLat is attenuated (Fig 4D bottom panel). Figure 4C and D confirmed that burst duration (3, 5 and 7 GC stimulations) strongly affect PC behavior.

To further study the effect of synaptic parameters, we measured spike gains in 12 different combinations of E and I synaptic strengths and two different E facilitation for each ΔLat (i.e., 24 different synaptic strength combinations per E/I delay and frequency). To evaluate these combinations in a simple way we rendered the results by monitoring ΔLat influence at 2 burst durations (3 and 7 stimulations), at which the switch between a net inhibition to a net excitation was observed (one example for the set of parameters used in Figure 4 is shown in Figure 5A and for all combinations at 40 Hz in Figure 5B). As shown in Figure 4C, varying ΔLat from negative to positive values increased spike gains, hence trajectories always move toward the upper right part of the plot. At 40 Hz (Figure 5B), the length of the trajectories (i.e., the total change in spike gain) is inversely proportional to the initial spike gain, and the strongest is the inhibition for 3 stimulations, the longer is the trajectory. Therefore, we converted each trajectory into a cumulative Euclidian distance in the 2-dimensional space spanned by spike gain for stimulus with 3 and 7 spikes (i.e., the total change in spike gain, see Methods and Figure 5A) and all 24 combinations were rendered for stimulation frequencies ranging from 20 to 200 Hz frequencies (**Figure 5C**). In this rendering, the longer the distance, the higher is the effect of ΔLat. As shown in Figure 4, mid-range frequencies (20-50 Hz) yield highest distance particularly for negative and low spike gains. These results clearly show that E/I-delays have a large effect on spike gain in PCs but only at low input frequencies. To better illustrate the influence of stimulation frequency on ΔLat effects, we estimated the distribution of distance at different frequencies (**Figure 5D**) and measured the entropy of the distribution (**Figure 5E**, see Methods). In this analysis the higher is the entropy the wider is the distribution and therefore the bigger is the effect of ΔLat. This revealed that ΔLat affect the spike gain at a wide range of frequencies but, again, their effect is the strongest at mid-range frequencies (20-75 Hz, **Figure 5D,E**). Taken together, when synapses show physiological STDs (Grangeray-Vilmint et al., 2018), a distribution of ΔLat as observed in the experimental dataset can expand the dynamic range of GC influence of PC discharge (spike gain) in biologically relevant range of frequencies.

**Figure 5:**
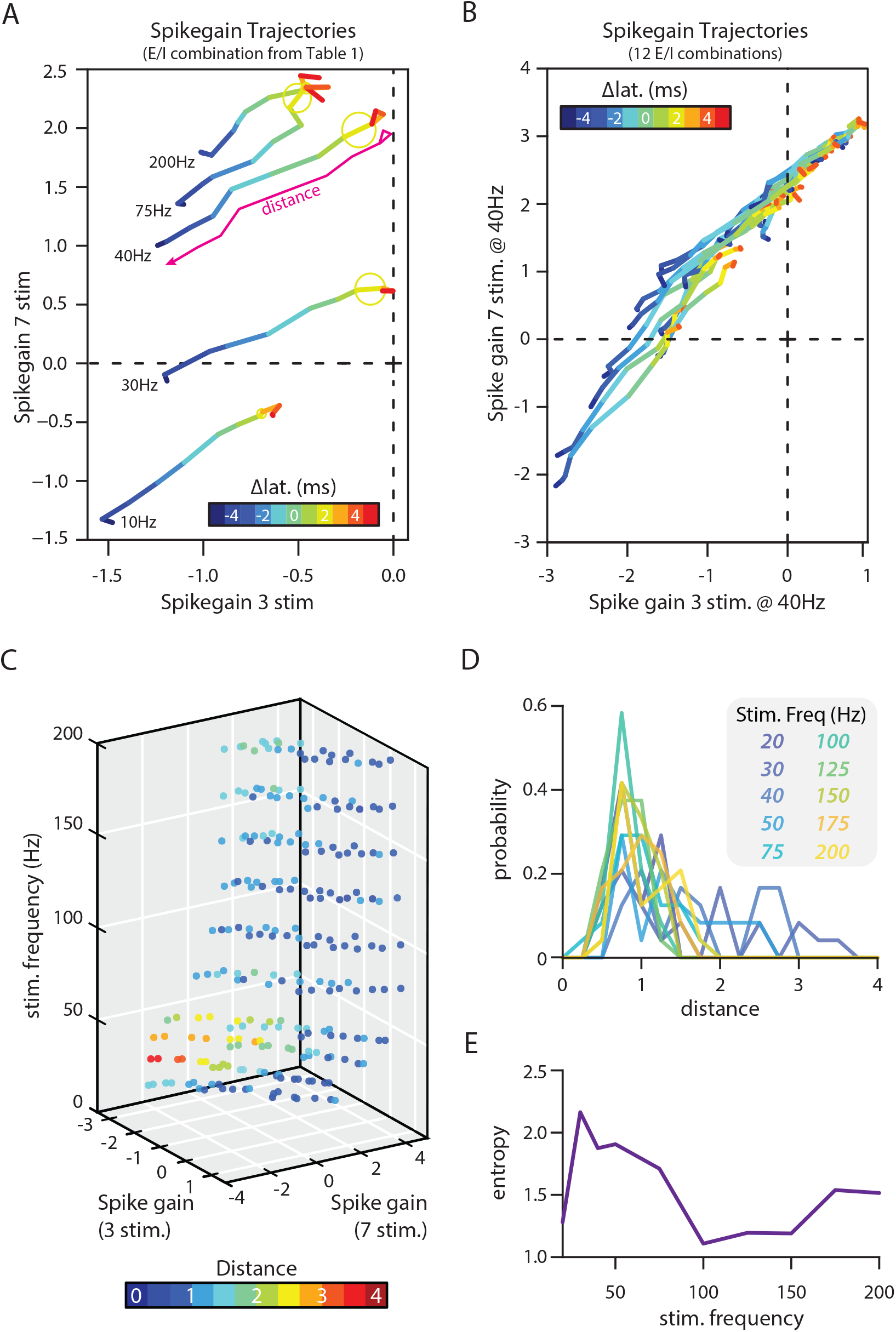
Variable ΔLat expand PC discharge range in response to biologically relevant GC inputs. **A**, Trajectories of PC spike gain for 3 and 7 stimulations when ΔLat vary from −5 ms (blue) to +5 ms (red) at various simulation frequencies (10 to 200Hz). STD are defined in table 1. The magenta line illustrates the distance measured at 40Hz (see methods). Yellow circles emphasize the trajectory measured in FFI (ΔLat=+2ms). **B**, Trajectories of PC spike gain for 3 and 7 stimulations when ΔLat vary from −5 ms (blue) to +5 ms (red) at 40 Hz stimulation frequency. Each line represents one combination of E, I and STD (12 combinations, see text and Table 1). **C**, Trajectories shown in B transformed into Euclidian distances (i.e., blue: no influence of E/I delay to red: strong influence from E/I delay, see methods) and calculated for stimulation frequency from 10 Hz to 200 Hz. EI delays largely contribute to modulation of PC discharge in case of low initial spike gain. **D**, distribution of distance for each combination of E, I and STD at different frequencies. **E**, entropy of distributions shown in D at varying frequencies.

## Discussion

Here we have shown that broad MF or single rosette stimulations in the cerebellar cortex leads to diverse sequences of synaptic excitation and inhibition in PCs. While synaptic strengths appeared correlated within these sequences, the delay between E and I ranged from negative to positive values extending the possibilities for a given set of synaptic inputs to influence PC discharge. While it seems easy to predict the modulation of PCs firing rate in response to nominal synaptic inputs *a priori* (i.e., GC input = acceleration and/or MLI input = deceleration), our experimental results supported by further simulations suggest that it remains crucial to consider the subsequent timing in which such inputs target PCs. Since the architecture of the MF-GC-MLI-PC pathway inherently leads to diverse E/I sequences (as summarized in Figure 1A), we therefore propose that the selective combination of STDs at GC-PC and GC-MLI-PC synapses allows PCs to distinguish between multiple synaptic inputs based on their temporal occurrence.

### Single or surface illumination evoked temporal E/I decorrelation in the MF-GC-(MLI)-PC pathway

In this study, we addressed the temporal aspect of E and I integration of the MF-GC-MLI-PC pathway in PCs by using single rosette and surface illumination protocols. Single rosette illumination elicited a single action potential and a steady depolarization allowing for a precise onset of vesicular release at the MF-GC synapses. We postulate that such direct depolarization, while not physiological, ensured a fast and reliable transmission by releasing most of the synaptic vesicles of the ready releasable pool (Jackman et al., 2014). Despite this direct illumination, delays between E and I in PCs were not only highly variable, but also incompatible with classical FFI during which GCs excite both PC and MLIs. ΔLat, particularly negative ΔLat, can only be accounted for by MF targeting different groups of GCs, indicating that the illumination leads to an antidromic propagation of the action potential elicited in the rosette in all the branches of the MF in the GC layer. Indeed, this behavior mimics physiological properties as incoming action potentials from the pre-cerebellar nuclei likely invade all the MF collaterals in the cerebellum. Anatomical reconstructions showed that a single MF gives rise to many distant rosettes in a cerebellar lobule (Shinoda et al., 2000; Sultan, 2001; Quy et al., 2011; Na et al., 2019), activating discrete and distinct GC clusters that may target the same PCs as shown previously (Valera et al., 2016; Spaeth et al., 2022). This latter configuration is also mimicked by the surface illumination used in our experiments which showed similar results than single rosette excitation but with a wider range of ΔLat. These findings demonstrate that MFs contact GCs that later target MLIs or PCs independently. Beyond ΔLat, we also observed that MF illumination can lead to pure inhibitory synaptic transmission, in agreement with the non-overlapping GC-PC and GC-MLI connectivity maps described in Valera et al (2016). In this latter study, by recording either PCs or MLIs at a given location in the cerebellar cortex, we described discrete areas of the GC layer that preferentially connect MLIs or PCs predicting that GC may lead to only inhibitory inputs to PCs. Altogether, in combination with the temporal expansion observed at the MF-GC connection (Chabrol et al., 2015), our data predict that E and I can be temporally de-correlated at the GC-PC connection.

In an FFI circuit when excitatory synapses are facilitatory and inhibitory synapses are depressing, it is optimal to represent input activity as a sparse code in which only a few neurons carry the stimulated-related information (Tauffer and Kumar, 2021). In such a scenario, stimulus related activity of PCs will be correlated, even if we assume a distribution of synaptic weights. However, if each GC projection on to a PC is associated with a slightly different delay, PC responses can be decorrelated. So besides, controlling the sign and magnitude of the PC responses, ΔLat distribution can also contribute to decorrelate PC responses.

Based on these results, it can be argued that E and I synaptic weights would be de-correlated as well. However, we showed that E and I synaptic weights recorded in PCs are correlated (Figure 3) as already shown in the cerebral cortex between parvalbumin interneurons and pyramidal cells (Xue et al., 2014). Equalized E/I ratio in the cerebral cortex is activity-dependent and spatially regulated independently at individual inhibitory synapses. Therefore, we suggest that similar rules may apply in the cerebellar GC-MLI-PC FFI pathway and individual GC clusters might be selected by PCs to ensure proper E/I balance in PCs. These findings are in agreement with the spatial organization of the GC-PC connectivity maps that correlate with specific locomotor context at the level of individuals (Spaeth et al., 2022). FFI circuits similar to the GC-MLI-PC with synapses that show STD are ubiquitous in the brain. Therefore, these results and insights are likely to be applicable to other brain regions.

### Extending temporal processing in the MF-GC-MLI-PC pathway

In the cerebellum, most principal neurons are spontaneously active (PC, nuclear cells) and MF inputs constantly convey sensorimotor information at a wide range of frequency to the GC layer. Typically at frequencies above 20 Hz, STD modify vesicular release (Perkel et al., 1990; Llano et al., 1991; Atluri and Regehr, 1996; Isope and Barbour, 2002; Valera et al., 2012; Doussau et al., 2017; Grangeray-Vilmint et al., 2018) and modifies E/I balance on a spike by spike basis. In this study, we extend our previous results showing that short term synaptic plasticity in the GC-MLI-PC pathway influences PC discharge (Grangeray-Vilmint et al., 2018). In particular, we show that E/I balance and PC discharge are also controlled by ΔLat in a frequency dependent manner. We notably identified that ΔLat have a strong effect on E/I balance and PC discharge only when input is occurring in a mid-range frequency (20-50 Hz; Figure 4 and 5). Indeed, MLIs fire at medium range frequencies in vivo and are synchronized together via gap junctions (Kim and Augustine, 2021). Many studies showed that MLI discharge is temporally controlled by electrical and chemical synapses (Mann-Metzer and Yarom, 1999; Alcami and Marty, 2013; Kim et al., 2014; Rieubland et al., 2014; Hoehne et al., 2020). Spikelet propagation in the MLI network ensure synchronization, while chemical synapses curtail integration time window (Hoehne et al., 2020). Altogether, selecting groups of GCs targeting specifically MLIs or PCs or both in combination (Valera et al., 2016; Spaeth et al., 2022) with specific short term dynamics (Chabrol et al., 2015; Dorgans et al., 2019) can enhance both the dynamic range and the resolution for temporal encoding in the MF-GC-PC pathway. Furthermore, we and others demonstrated that long term plasticity (LTP and LTD) at the GC-PC synapses is gated by MLIs (Binda et al., 2016; Rowan et al., 2018). We then postulate that the independent regulation of GC-MLI-PC synapses can influence plasticity induction from specific cluster of GCs. These features may underlie cerebellar module communication and motor coordination (Apps et al, 2018). Finally, an appealing hypothesis would be that temporal dynamics between 2 different groups of MF inputs (e.g. from different modalities) might lead to the partial or the total suppression of one input. These properties might account for one of the main roles of the cerebellum, which is to suppress the expected sensorimotor feedback in order to process further the unexpected inputs.

## Materials and Methods

All experiments were conducted in accordance with the guidelines of the Ministère de l’Education Supérieure et de la Recherche and the local ethical committee, the Comité Régional En Matière d’Expérimentation Animale de Strasbourg (CREMEAS) under the agreement delivered to the animal facility Chronobiotron (UMS3415, University of Strasbourg)

### Pre-cerebellar nuclei rAAVs-mediated transduction

In vivo stereotaxic injections of rAVVs viral particles were performed as previously described (Valera et al. 2016). CD1 male mice (P 21) were anesthetized by a brief exposure to isoflurane 4 % and anesthesia was maintained by intraperitoneal injections of a mixture of ketamine (100 mg/kg), medetomidine (1 mg/kg) and acepromazine (3 mg/kg). rAAVs 9/2 particles carrying the cDNA for ChRd2(H134R)-YFP under the hSyn promoter (3.38 10^13^ GC/ml; Penn Vector Core, Pennsylvania) were unilaterally injected in the cuneate nucleus at an approximate speed of 250 nl/min via a graduated pipette equipped with a piston for manual injections. A final volume of 1.5 μl was delivered by two injections (0.75 μl/injection) separated by 0.2 mm in the antero-posterior direction; after that half of the virus volume was delivered, the pipette was raised up 0.2 mm and maintained in place until the end of the injection. For effective virus diffusion, the pipette was left in place at least 5 minutes following injection. Injections coordinates were determined from The Mouse Brain Atlas (Franklin and Paxinos, 2007) and corrections based on tissue markers were applied to counterbalance the variability of the CD1 outbred background (from lambda, starting point: AP 2.73 ± 0.14, Lat 1.36 ± 0.08, DV: 5.1 ± 0.09, mean ± SEM). At the end of the injection, antipamezole (1 mg/kg) were administered to the mice via intraperitoneal injection to favor recovery from anesthesia.

### Slice preparation

Acute cerebellar transverse slices were prepared from injected mice three weeks after injection. Mice were anesthetized by a brief exposure to isoflurane 4 %, decapitated and the cerebellum rapidly dissected in ice-cold artificial cerebrospinal fluid (aCSF) bubbled with carbogen (95% O_2_, 5% CO_2_) and containing (in mM): NaCl 120, KCl 3, NaHCO_3_ 26, NaH_2_PO_4_ 1.25, CaCl_2_ 2, MgCl_2_ 1 and glucose 16. 300 μm-thick transverse slices were cut (Microm HM 650V, Microm, Germany) in ice-cold sucrose-based cutting solution bubbled with carbogen (95% O_2_, 5% CO_2_) and containing (in mM): sucrose 246, KCl 4, NaHCO_3_ 26, CaCl_2_ 1, MgCl_2_ 5, glucose 10 and kynurenic acid 1. Slices were then transferred in bubbled aCSF at 34°C and they were allowed to recovery for at least 30 minutes before starting experiments. Following recovery, slices were maintained at room temperature for the rest of the day.

### Electrophysiology

Electrophysiology experiments were performed at room temperature in a recording chamber continuously perfused with bubbled aCSF supplemented with (in mM): strychnine 0.001, D-APV 0.05, DPCPX 0.0005. AM251 0.001, CGP52432 0.001 and JNJ16259685 0.002. To allow the full illumination of the surface, slices were rapidly mounted on glass coverslips coated with poly-L-lysine (1 mg/ml) right before transferring them to the recording chamber.

Patch clamp pipettes were pulled from borosilicate glass to a final resistance of 3-4 MΩ when filled with the following solution (in mM): cesium methanesulfonate 135, NaCl 6, MgCl_2_ 1, HEPES 10, MgATP 4, Na2GTP 0.4, EGTA 1.5, QX314Cl 5, adjusted to pH = 7.3. Whole-cell patch clamp recordings were performed with a Multiclamp 700B amplifier (Molecular Devices, USA) and acquired with WinWCP 4.2.1 freeware (John Dempster, SIPBS, University of Strathclyde, UK); whole-cell currents from PCs were filtered at 2 kHz and digitized at 50 kHz; series resistance was monitored during the experiments and compensated by 70-80%.

Loose cell-attached recordings were obtained using a Multiclamp 700B amplifier (Molecular Devices, USA) and acquired with WinWCP 4.2.1 freeware (John Dempster, SIPBS, University of Strathclyde, UK). Rosettes were recorded with 5 MΩ glass pipettes (borosilicate) and potential was held at 0mV for all recordings. The internal pipette solution contained (in mM): NaCl 120, KCl 3, HEPES 10, NaH_2_PO_4_ 1.25, CaCl_2_ 2, MgCl_2_ 1 and glucose 10 (SigmaAldrich, USA). Osmolarity and pH were respectively set at 295 mOsm and 7.3. Recordings were lowpass filtered at 2.6 kHz then sampled at 20-50 kHz. All experiments were performed at room temperature (23°C) using the same bubbled aCSF than for slices preparation. For controls, NBQX (20 μM, Sigma-Aldrich, USA) and lidocaine (1mM, Sigma-Aldrich, USA) were bath-applied respectively to block AMPA receptor transmission and action potential propagation.

For GC stimulation, a glass pipette (borosilicate) containing (in mM) : NaCl 120, KCl 3, HEPES 10, NaH_2_PO_4_ 1.25, CaCl_2_ 2, MgCl_2_ 1 and glucose 10 (SigmaAldrich, USA) was inserted in the deepness of the GC layer. GCs were electrically stimulated (3-5μA, 100μs) using a constant current delivered by an isolated stimulator (model DS3, Digitimer Ltd, England).

### Photostimulation

Surface mossy fibers blue light-mediated activation was obtained by a two dimension-scan mode of a confocal microscope (FV300, Olympus, Japan) equipped with a diode-pumped solid-state blue laser (473 nm, CrystalLaser, USA); stimulation (50 ms blue light pulses through a 20X objective, 83×83 μm scan area) was driven by a Programmable Acquisition Protocol Processor (PAPP, Fluoview 300) at successive positions along the longitudinal axis. The stimulation protocol was repeated between 3 and 7 times and average traces used for analysis. MFs-dependent EPSCs and IPSCs were isolated by clamping Purkinje cells at – 60 mV and 0 mV respectively.

Unitary stimulations for single illumination protocol and loose-cell recordings of MF terminals were performed using a 460nm blue LED (Prizmatix, Israel). The illuminated area was restricted to a single rosette using a DMD-array (Mosaïc, Andor Technology, Northern Ireland) through a 40x objective (Olympus, Japan). Steady illumination power was set at 2-4mW/mm^2^. The stimulation protocol was repeated from 10 to 20 times and average traces were used for analysis.

### Simulation

We used a previous model of GC-MLI-PC pathway that we published in Grangeray-Vilmint et al. (2018) to study the effect of E/I delay and input frequency on the PC output. The model was simulated using the NEural Simulation Tool (NEST) environment (http://www.nestsimulator.org/).

#### PC neuron

The Purkinje cell was modelled as a conductance-based point neuron. The membrane voltage dynamics of the PC was given by:

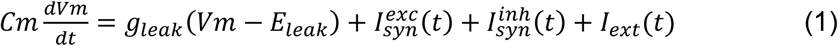

where *Cm* is the membrane capacitance, *g*_*leak*_ is the leak conductance, *E*_*leak*_ is the resting potential of the neuron. When membrane potential reached the spiking threshold (−55mV) a spike was elicited and the membrane was clamped at the resting membrane potential for 2 ms (refractory period). 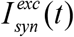 represents the total excitatory inputs arriving from the GrCs and 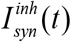 represents the total inhibitory inputs arriving from the inhibitory interneurons. In the absence of any specific stimulus PC spike at about ≈ 30 Hz in a quasi-periodic manner (*CV*_*isi*_ ≈ 0.2). To mimic this ongoing activity in our simple point neuron model of the PC we injected noisy excitatory current, *I*_*ext*_ (*t*), corresponding to a spike train drawn from a sinusoidally modulated Γ-process (frequency =37 Hz; order = 4). The magnitude of the sinusoidal modulation was 10% of the average input rate which was set to 2000 spikes/sec to obtain 30 spikes/sec as the output firing rate in the model neuron. For each trial we used a different realization of the sinusoidally modulated Γ-process to generate the background activity in the PC. Despite being an over simplification of the dynamics of PCs, this simple model allows us to study the effect of synaptic dynamics on the integration of external input by a FFI circuit motif. Complex non-linear dynamics of PCs could have obscured these synaptic effects.

#### Synapse model

Synapses were modeled as a conductance transient

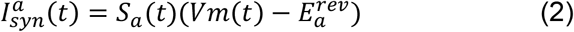

where *S*_*a*_*(t)* (*a* ∈ {exc., inh.}) is the synaptic transient, *V*_*m*_ is the membrane potential and 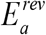 is the reversal potential of the synapse. See Table 2 for the parameters. Each presynaptic spike induced an alpha function shaped conductance transient in the postsynaptic PC:

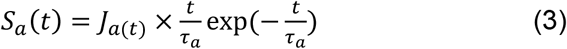

where τ_a_ is the synaptic time constant and *J*_*a*_*(t)* is the amplitude of the synapse. When synapses are static *J*_*a*_*(t)* is constant and when synapses change according to the history of the previous presynaptic spikes, the temporal evolution of *J*_*a*_*(t)* is given by the eq. 8.

#### Short-term dynamics of synapses

To model the STD of the PSP amplitude we used a deterministic model proposed by Tsodyks et al. (Tsodyks and Markram, 1997).

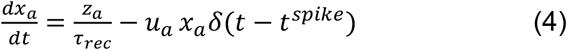

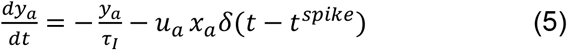

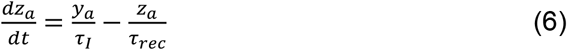

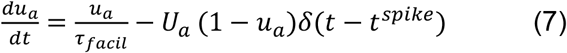

where *x*_*a*_, *y*_*a*_, *z*_*a*_ (*a* ∈ {exc., inh.}) are the 3 state variables that describe the state dependent use of the synaptic resources in the recovered (*x*_*a*_), active (*y*_*a*_), and inactive (*z*_*a*_) states, respectively. *t*^*spike*^ is the timing of the previous spike. τ_*I*_ is the decay time constant of the postsynaptic current, *τ*_*rec*_ is the time it takes the synaptic resources to recover from their inactive state. The variable *u*_*a*_*(t)* keeps track of the use of the synaptic resources at a synapse. To model synaptic facilitation the variable *u*_*a*_ is instantaneously increased by a small amount *U*_*a*_ at the time of spike and then returns to the baseline value with a time constant *τ*_*facil*_ (eq. 7).

The effective synaptic weight was given by,

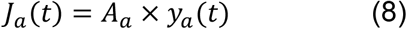

To systematically change the nature of the synapses from facilitating to depressing, we changed the variable *U*_*a*_ while keeping all other variables unchanged. Because we kept all the time constants unchanged in the model, going from facilitation to depression did not affect the frequency dependence of the synapses. The parameter *A*_*a*_ was changed to increase the amplitude of the postsynaptic potential independent of the STD. The values of these various parameters are provided in the Table 1.

### Data Analysis

Electrophysiological recordings were analyzed with custom scripts and routines written in Python 3.9 using the following packages: Pandas 1.3, Scipy 1.6, NumPy 1.19, and Neo 0.9. Complete code is available at https://github.com/ludo67100/MFDeltaLat.

Both synaptic charges and latencies were computed on averaged signals (3 to 20 stimulations). The latency was defined as the time difference between stimulation onset and the intersection between a linear fit of the current rising phase and the baseline of the signal (see examples in Figure 2E). ΔLat were computed by subtracting IPSC latency from EPSC latency in each experiment. Synaptic charges were measured as the integral of the signal in a 100ms time window starting from the onset of light stimulation.

Statistical tests were performed with corresponding packages from scipy.stats module. Alternative and method argument were respectively set to ‘two-sided’ and ‘auto’ unless otherwise reported. P-values <0.05 are indicated in figures. For the ΔLat measurement following direct, electrical GC stimulation, the dataset recorded for this experiment has been combined with the dataset from Grangeray-Vilmint et al. (2018).

Spike gain: we binned the spiking activity across all the 200 trials to obtain peri-stimulus-time-histogram (PSTH: bin width = 2ms). Next, we measure the mean of the PSTH in the prestimulus duration (i.e. in the interval 50-200ms). Next, we calculated spike gain as mean and sum of the normalized PSTH (Figure 4A).

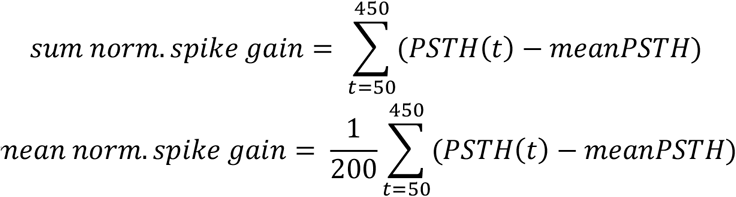

We normalize with 200 as we have 200 bins of 2ms each. In the Fig. 4 we have plotted *mean norm, spike gain* but the results are same if we plot *sum norm. spike gain*. For rendering the Fig. 4B we further normalized these spike gain measures with respect to the spike gain measured as E/I delay of +2ms.

Spike grain trajectory distance: We have two spike gains, one for the 3 stim and one for 7 stim. We can represent the neuron output as a point in a two-dimensional space spanned by spike gain for 3 stim (x-axis) and 7 stim (y-axis) See Fig. 5A). To quantify the effect of E/I spike latency we measured Euclidian distance between neuron output for two different E-I latency:

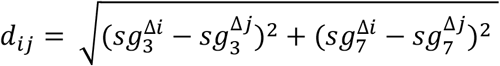

Where 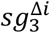 is the spike gain for 3 stim case for *i*th E/I-latency and 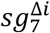 is the spike gain for 7 stim case for *i*th E/I-latency. As we varied E/I delay from −5ms to +6ms we had 11 possible distances for each increment in the EI delay. To quantify the total effect of the full range of EI-latency we define cumulative distance as

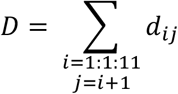

#### Entropy of the distances

Let *d*_*ij*_*(f)* where *i = 1:1:11 and j = i+*1 is the set of all distances for a specific frequency (see above). We first estimated the count histogram of all *d*_*ij*_ *(f)*. This histogram was normalized to have a unit area. Now we treated this histogram as a probability distribution and estimated the entropy using Shannon’s formula

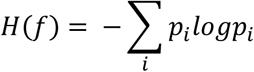

Where *p*_*i*_ is the normalized count of a specific distance in the set *d*_*ij*_*(f)*. The sum runs over all distances. Higher *H(f)* indicates a wider distribution and when all distances are identical, *H(f)=*0.

## Acknowledgments

This work was supported by the Centre National pour la Recherche Scientifique (CNRS), the Université de Strasbourg, the Agence Nationale pour la Recherche (ANR-2015-CeMod, ANR-2019-MultiMod, ANR-2019-NetOnTime) and by the Fondation pour la Recherche Medicale to PI (# DEQ20140329514). LS was funded by fellowships from the Ministère de la Recherche and Fondation pour la Recherche Médicale (#FDT201805005172). AK was funded by Swedish Research Council (2018-03118), StratNeuro, Digital Future and University of Strasbourg Institute for Advanced Studies (USIAS) fellowship. We thank Dr. Sophie Reibel-Foisset, Dominique Ciocca and the staff of the animal facility (Chronobiotron, UMS 3415 CNRS and Strasbourg University) for technical assistance. We thank Dr Bernard Poulain, Dr Antoine Valera, Dr Matilde Cordero-Erausquin, Dr Didier Desaintjan and Dr Frédéric Doussau for critical discussions.

## References

Alcami P, Marty A (2013) Estimating functional connectivity in an electrically coupled interneuron network. Proc Natl Acad Sci U S A 110:E4798–807

Apps R et al. (2018) Cerebellar Modules and Their Role as Operational Cerebellar Processing Units: A Consensus paper [corrected]. Cerebellum 17:654–682

Apps R, Hawkes R (2009) Cerebellar cortical organization: a one-map hypothesis. Nat Rev Neurosci 10:670–681

Arenz A, Silver RA, Schaefer AT, Margrie TW (2008) The contribution of single synapses to sensory representation in vivo. Science 321:977–980

Atluri PP, Regehr WG (1996) Determinants of the time course of facilitation at the granule cell to Purkinje cell synapse. J Neurosci 16:5661–5671

Barbour B (1993) Synaptic currents evoked in Purkinje cells by stimulating individual granule cells. Neuron 11:759–769.

Binda F, Dorgans K, Reibel S, Sakimura K, Kano M, Poulain B, Isope P (2016) Inhibition promotes long-term potentiation at cerebellar excitatory synapses. Sci Rep 6:33561

Brown ST, Raman IM (2018) Sensorimotor Integration and Amplification of Reflexive Whisking by Well-Timed Spiking in the Cerebellar Corticonuclear Circuit. Neuron 99:564–575.

Brunel N, Hakim V, Isope P, Nadal J-P, Barbour B (2004) Optimal information storage and the distribution of synaptic weights: perceptron versus Purkinje cell. Neuron 43:745–757

Chabrol FP, Arenz A, Wiechert MT, Margrie TW, DiGregorio DA (2015) Synaptic diversity enables temporal coding of coincident multisensory inputs in single neurons. Nat Neurosci 18:718–727

Chadderton P, Margrie TW, Häusser M (2004) Integration of quanta in cerebellar granule cells during sensory processing. Nature 428:856–860

Dorgans K, Demais V, Bailly Y, Poulain B, Isope P, Doussau F (2019) Short-term plasticity at cerebellar granule cell to molecular layer interneuron synapses expands information processing. Elife 8:1–24

Doussau F, Schmidt H, Dorgans K, Valera AM, Poulain B, Isope P (2017) Frequency-dependent mobilization of heterogeneous pools of synaptic vesicles shapes presynaptic plasticity. Elife 6:e28935

Eccles JC, Ito M, Szentagotai J (1967) The cerebellum as neuronal machine. Berlin: Springer-verlag.

Franklin K, Paxinos G (2007) The Mouse Brain in Stereotaxic coordinates, 3rd ed. Academic Press, San Diego.

Grangeray-Vilmint A, Valera AM, Kumar A, Isope P (2018) Short-Term Plasticity Combines with Excitation-Inhibition Balance to Expand Cerebellar Purkinje Cell Dynamic Range. J Neurosci 38:5153–5167

Hoehne A, McFadden MH, DiGregorio DA (2020) Feed-forward recruitment of electrical synapses enhances synchronous spiking in the mouse cerebellar cortex. Elife 9:e57344

Isaacson JS, Scanziani M (2011) How inhibition shapes cortical activity. Neuron 72:231–243

Isope P, Barbour B (2002) Properties of Unitary Granule {Cell→Purkinje} Cell Synapses in Adult Rat Cerebellar Slices. J Neurosci 22:9668–9678

Jackman SL, Beneduce BM, Drew IR, Regehr WG (2014) Achieving high-frequency optical control of synaptic transmission. J Neurosci 34:7704–7714

Jelitai M, Puggioni P, Ishikawa T, Rinaldi A, Duguid I (2016) Dendritic excitation-inhibition balance shapes cerebellar output during motor behaviour. Nat Commun 7:13722

Jörntell H, Bengtsson F, Schonewille M, De Zeeuw CI (2010) Cerebellar molecular layer interneurons - computational properties and roles in learning. Trends Neurosci 33:524– 532

Jörntell H, Ekerot C-F (2006) Properties of somatosensory synaptic integration in cerebellar granule cells in vivo. J Neurosci 26:11786–11797

Kim J, Augustine GJ (2021) Molecular Layer Interneurons: Key Elements of Cerebellar Network Computation and Behavior. Neuroscience 462:22–35

Kim J, Lee S, Tsuda S, Zhang X, Asrican B, Gloss B, Feng G, Augustine GJ (2014) Optogenetic mapping of cerebellar inhibitory circuitry reveals spatially biased coordination of interneurons via electrical synapses. Cell Rep 7:1601–1613

Llano I, Marty A, Armstrong CM, Konnerth A (1991) Synaptic- and agonist-induced excitatory currents of Purkinje cells in rat cerebellar slices. J Physiol 434:183–213

Mann-Metzer P, Yarom Y (1999) Electrotonic coupling interacts with intrinsic properties to generate synchronized activity in cerebellar networks of inhibitory interneurons. J Neurosci 19:3298–3306

Mittmann W, Koch U, Häusser M (2005) Feed-forward inhibition shapes the spike output of cerebellar Purkinje cells. J Physiol 563:369–378

Na J, Sugihara I, Shinoda Y (2019) The entire trajectories of single pontocerebellar axons and their lobular and longitudinal terminal distribution patterns in multiple aldolase C-positive compartments of the rat cerebellar cortex. J Comp Neurol 527:2488–2511

Palay SL, Chan-Palay V (1974) Cerebellar Cortex. Berlin, Heidelberg: Springer Berlin Heidelberg.

Perkel DJ, Hestrin S, Sah P, Nicoll RA (1990) Excitatory synaptic currents in Purkinje cells. Proc Biol Sci 241:116–121

Quy PN, Fujita H, Sakamoto Y, Na J, Sugihara I (2011) Projection patterns of single mossy fiber axons originating from the dorsal column nuclei mapped on the aldolase C compartments in the rat cerebellar cortex. J Comp Neurol 519:874–899

Rieubland S, Roth A, Häusser M (2014) Structured connectivity in cerebellar inhibitory networks. Neuron 81:913–929

Rowan MJM, Bonnan A, Zhang K, Amat SB, Kikuchi C, Taniguchi H, Augustine GJ, Christie JM (2018) Graded Control of Climbing-Fiber-Mediated Plasticity and Learning by Inhibition in the Cerebellum. Neuron 99:999–1015.

Santamaria F, Tripp PG, Bower JM (2007) Feedforward inhibition controls the spread of granule cell-induced Purkinje cell activity in the cerebellar cortex. J Neurophysiol 97:248–263

Shinoda Y, Sugihara I, Wu HS, Sugiuchi Y (2000) The entire trajectory of single climbing and mossy fibers in the cerebellar nuclei and cortex. Prog Brain Res 124:173–186

Spaeth L, Bahuguna J, Gagneux T, Dorgans K, Sugihara I, Poulain B, Battaglia D, Isope P (2022) Cerebellar connectivity maps embody individual adaptive behavior in mice. Nat Commun 13:580

Sultan F (2001) Distribution of mossy fibre rosettes in the cerebellum of cat and mice: evidence for a parasagittal organization at the single fibre level. Eur J Neurosci 13:2123–2130

Tauffer L, Kumar A (2021) Short-Term Synaptic Plasticity Makes Neurons Sensitive to the Distribution of Presynaptic Population Firing Rates. eNeuro 8(2)

Tsodyks M V, Markram H (1997) The neural code between neocortical pyramidal neurons depends on neurotransmitter release probability. Proc Natl Acad Sci U S A 94:719–723

Valera AM, Binda F, Pawlowski SA, Dupont J-L, Casella J-F, Rothstein JD, Poulain B, Isope P (2016) Stereotyped spatial patterns of functional synaptic connectivity in the cerebellar cortex. Elife 5:e09862

Valera AM, Doussau F, Poulain B, Barbour B, Isope P (2012) Adaptation of Granule Cell to Purkinje Cell Synapses to High-Frequency Transmission. J Neurosci 32:3267–3280

Vincent P, Marty A (1996) Fluctuations of inhibitory postsynaptic currents in Purkinje cells from rat cerebellar slices. J Physiol 494 (Pt 1):183–199

Xue M, Atallah B V., Scanziani M (2014) Equalizing excitation-inhibition ratios across visual cortical neurons. Nature 511:596–600

